# Directly and indirectly determinable rate constants in Michaelian enzyme-catalyzed reactions

**DOI:** 10.1101/2022.12.27.522020

**Authors:** Ikechukwu Iloh Udema

## Abstract

Backed by kinetic schemes, attempts had been made to derive equations for the calculation of all zero-order first-order rate constants (ZOFORC) for the activation of the enzyme-substrate (ES) complex and its deactivation, *k*_+2_ and *k* _−2_, respectively. The values of ZOFORC, including the kind for the dissociation of the enzyme-product complex (EP) to free enzyme (E) and product (P), are hardly reported. The methods of research were primarily Bernfeld and Lineweaver methods. The goal of the research was to determine ways for the utilization of experimental data for the determination of verifiable and quantifiable rate constants, with the following objectives: 1) To derive equations for the first order rate constants, *k*_+2_ and *k*_*-*2_, for the activation of ES and its deactivation, respectively; 2) to quantify by calculation the first order rate constant for product release; 3) ultimately quantify the rate constants, *k*_-2_ and *k*_+2_; and 4) to advise the reactor, process, chemical engineers, *etc*. in different industrial concerns on the usefulness of *k*_-2_ and *k*_+2_. The value of ZOFORC for the dissociation of EP to free E and P is 3.155 exp. (+5)/min; the values of *k*_+2_ and *k*_-2_ are 3.513 exp. (+4) and 2.377 exp. (+8)/min, respectively. Ultimately, it is imperative for all stakeholder groups to devise means of controlling the enzymatic rate of catalysis by manipulating the magnitudes of *k*_+2_ and *k*_-2_ in particular. The derived equations can be fitted to the experimentally generated and calculated data. A future research project should entail conducting the assay under optimum conditions so as to verify possible variations in the ZOFORC values when compared with values generated outside optimum conditions.

## 1. Introduction

When one examines the literature with a strong interest in the rates of reactions catalyzed by enzymes, one comes up with the finding that there are a lot of kinetic schemes with which to generate kinetic equations, rates, and rate constants; the most popular and simplest of such schemes is: E + S ⇌ES → E + P, where E, S, ES, and P are the symbols for the enzyme, the substrate, the enzyme-substrate complex, and the product, respectively. The scheme has been described as being hypothetical [1]. However, the scheme illustrates the formation of an enzyme-substrate complex, without which progress towards product formation would be impossible. As stated elsewhere [2], the earliest known mechanism of enzyme action (or function) is anchored on the “induced fit” hypothesis or model of Daniel Koshland, Jr. [3] and the “lock and key hypothesis” of Emil Fischer [4]. As a result, it appears that any discussion and concern about the rates of enzyme-catalyzed reactions cannot be complete without mentioning the mechanism of binding. Incidentally, the induced-fit model seemed to have a vague meaning, leading to a more acceptable enzyme’s conformational transition (change) model [1]. The conformational change of this enzyme during encounter and interaction with substrate is thought to be the one that determines the specificity steps [1]. This notwithstanding, an earlier opinion is that the induced fit model seems to be gaining ground with the advent of the “conformation selection” principle [5–10], which postulates that all of the potential conformations of a given protein preexist and that once the ligand selects the most favored conformation, induced fit occurs and conformational change takes place [1].

The generation of rate equations and rate constants requires kinetic schemes far more complex than the generalized simple case given earlier. On the question of the complexity of kinetic schemes that can instill fear in undergraduates, one has the following to advise upon, as discussed elsewhere [11]: Every scheme generated needs to be explained in a stepwise manner. Still, there has been an exceedingly complex derivation of kinetic equations for rate and rate constants based on the rapid equilibrium assumption or combined assumptions of equilibrium and steady-state, where respectively, Michaelis and Menten [12] and Briggs and Haden [13] were the original contributors [14].

The paper by Strickland *et al*. [15] has as its goal the determination of dissociation constants and specific rate constants of enzyme-substrate complexes, achieved via two different schemes that depict two different mechanisms of enzymatic action; the equations derived based on the schemes, though simple, were not based on a detailed stepwise derivation. Simulations were largely explored for data generation and fitting the derived equations. In this research, a scheme formulated by Johnson [1] is explorable for the derivation of the equations for the first-order rate constants and the quantification of the kinetic constants that are not regularly featured in most enzymology studies. The scheme is thought to be very generalizable to hydrolytic enzymes, such as the amylolytic enzyme.

There is no doubt that the rate constants are very important from the standpoint of biological application, reactor design, drug detoxification and clearance, *etc*. But there is hardly any detailed explanation of how such equations were derived. There are also few attempts to fit the derived equations to experimental data for quantification by the calculation of multiple kinetic parameters. It is a well-documented fact in the literature [1, 11, 15, 16] that an enzyme-catalyzed reaction has different stages, viz., encounter complex formation, binding of substrate to the enzyme, conformational transformation to activated complex formation, bond breaking and making (the chemistry), and the release of product and free enzyme. This is possible if the activated complex formation is not aborted. If, for any undefined reason, it is suddenly aborted, the ES that is weakly bound together breaks down into free E and S. These issues have been highlighted in the literature [11].

Besides, the methods of study other than transient-phase kinetic studies using the stopped-flow method [18] for the determination of the first-order rate constants *k*_3_, for the process, EP → E + P (this needed update is for correctional purposes), *k*_ES_, for the process, E + S → ES, and *k*_-1_, for the process, ES → E + S, are given in the literature [19]. Nonetheless, backed by kinetic schemes, attempts had been made to derive equations for the calculation of all zero-order first-order rate constants (ZOFORC) for the activation of the enzyme-substrate (ES) complex and its deactivation, *k*_+2_ and *k*_*−*2_, respectively. The values of ZOFORC, including the kind for the dissociation of the enzyme-product complex (EP) to free enzyme (E) and product (P), are hardly reported. Considering the importance of rates in medicine, pharmaceutics, nutrition, engineering, *etc*., and in order to be able to utilize experimental data for the determination of verifiable and quantifiable rate constants as an important goal, it has become very imperative to conduct this research with the following objectives: 1) To derive equations for the first order rate constants, *k*_+2_ and *k*_*−*2_, for the processes, ES → E^#^S^#^ and E^#^S^#^ → ES, respectively; 2) to quantify by calculation the first order rate constant for product release; 3) ultimately quantify the rate constants, *k*_-2_ and *k*_+2_; and 4) to advise the reactor, process, chemical engineers, *etc*. in different industrial concerns on the usefulness of *k*_-2_ and *k*_+2_.

## 2. Theory

In this section, the kinetic schemes in the literature and selected kinetic equations are to be revisited, and new equations are to be derived and evaluated.

### 2.1 Examination of simple and practical kinetic schemes

There are a lot of kinetic schemes in the literature [1, 11, 16, 17]. The most reoccurring, as stated earlier, is the scheme given as: E+S⇌ES→E + P. However, the scheme that is more similar to the kind of scheme to be explored in this research is the kind given by Johnson [1]. This is scheme 1 below. In that scheme, the author was of the opinion that the rate constant (*k*_3_) for the process EP → E + P is much greater than *k*_2_, which was equated to *k*_cat_ (the catalytic rate, the turn-over number). The author has the impression that equilibrium exists between ES and EP. However, by indicating that the rate constant for the process EP → ES is equal to zero, there is an admission that such a process does not occur, if, in particular, the enzyme in this research is a hydrolase lacking the power to synthesize. These two views of the author are in line with the goal of this research, which entails showing that the overall duration of the enzyme-catalyzed reaction pathway is greater than the specific duration of each phase of the reaction pathway. Furthermore, enzymes primarily accelerate reactions by lowering the “energy barrier” via the formation of an activated enzyme-substrate complex (E^#^S^#^); this complex may exist for an infinitesimal duration, either proceeding to EP or ES if unfavorable conditions arise for whatever reason. Thus, in a scheme such as E + S ⇌ES ⇌E^#^S^#^ → EP→P, the rate constant for the process ES →E^#^S^#^ is designated as *k*_*+*2_, while the rate constant for the process E^#^S^#^→ES is designated as *k*_-2_. For the purpose of simplification, *k*_3_ may stand for the rate constant for the process E^#^S^#^ → EP→P that remains irreversible.

**Scheme 1:** Formation of enzyme-substrate complex (ES), transformation to enzyme-product complex (EP), and dissociation into free enzyme and product [1].

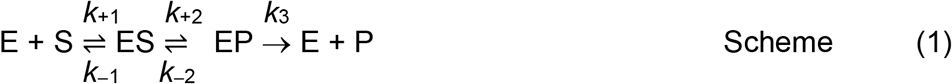

As explained in the literature [11], there could be an activated complex formation given as E^#^S^#^, which represents a transition intermediate between ES and EP. This is such that the likelihood of EP reversing to ES is impossible as long as the enzyme is neither a synthase nor a synthetase; however, there is a strong likelihood that E^#^S^#^ reverses back to ES for whatever reason. Assuming that E^#^S^#^ and EP have very similar life spans, there is a possibility of the process ES ⇌E^#^S^#^ → EP such that the real equilibrium lies between ES and E^#^S^#^. The second scheme, which summarizes the issues raised, is stated as follows:

Modified scheme 1, which shows the presence of an activated enzyme-substrate complex

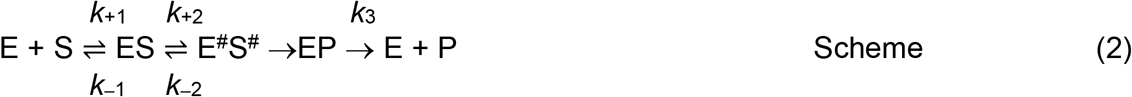

The complex, E^#^S^#^ has an infinitesimal life span, proceeding either to EP or ES; EP breaks up into E and P as quickly as it is formed. Therefore, the process depicted above is summarized as “E + S ⇌ES → E + P.” If the duration of the process (EP → E + P) is “*τ*”, the first order rate constant is denoted as *k*_3_ (this is = 1/*τ*). The equations (Eqs (1) and (2)) [1] to be revisited are given below.

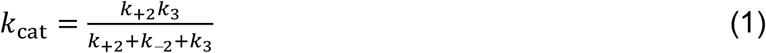

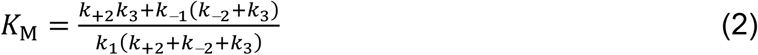

In order to ascertain the likely validity of the equations to be derived based on Eqs (1) and (2), both equations are examined. The outcome shows that both equations have a common denominator if Eq. (2) is recast as follows: From Eq. (1) is the following:

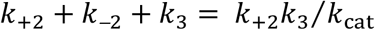

From Eq. (2) is the following:

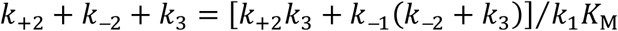

Therefore,

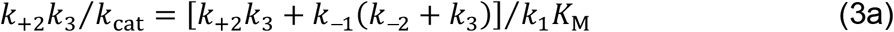

Making *k*_+2_ subject of the formula gives:

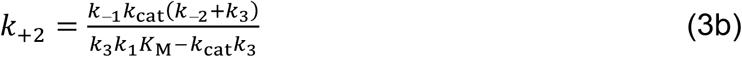

Making *k*_+2_ the subject of the formula in Eq. (1) yields:

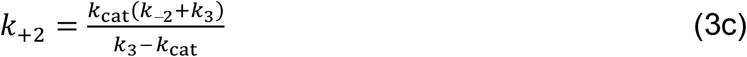

Hence, Eq. (3c) is the equivalent of Eq. (3b) such that:

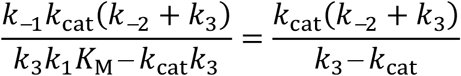

Making *K*_M_ subject of the preceding formula gives after simplification:

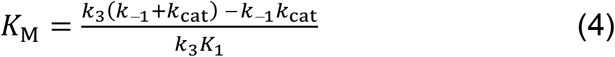

Meanwhile, *K*_M_*k*_1_ is = *k*_−1_ + *k*_cat_ such that the expansion of Eq. (4) gives first:

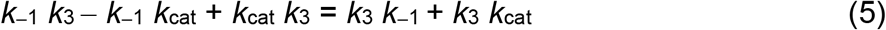

Elimination of common terms in the immediate preceding equation gives:

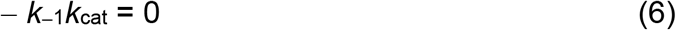

The result indicated by Eq. (6) shows that there may be a problem in either Eq. (1) or Eq. (2). Such conclusion may be too early at this stage.

However, Eqs (1) and (2) are revisited through a different route. From the two equations is given the following:

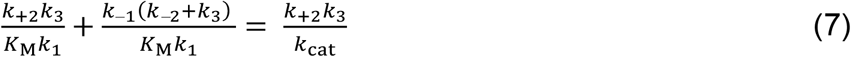

Factorizing and simplifying gives:

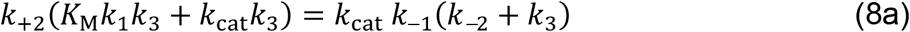

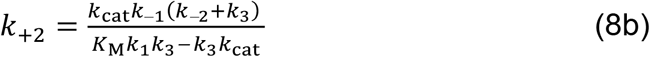

Directly from Eq. (1) is Eq. (3c) and bringing Eqs (3c) and Eq. (8b) together gives

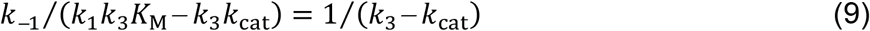

Realizing that *k*_1_*K*_M_ is = *k*_−1_ + *k*_cat_ and upon rearrangement of Eq. (9) one obtains:

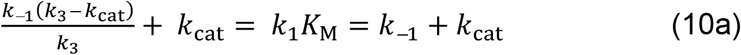

Equation (10a) leads to:

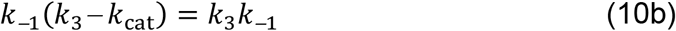

Equation (10b) has two possible outcomes:

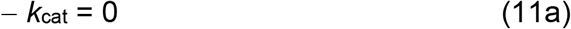

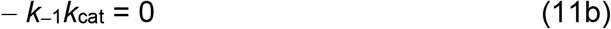

Equation (11b), which is a reproduction of Eq.(6) and Eq.(11a) gives enough evidence for the invalidity of either Eq. (1) or Eq. (2).

### 2.2 The derivation of the equations for the calculation of kinetic parameters that are not discernible in a reaction pathway

There is an argument that the data obtained in the steady-state provide only indirect information to define the pathway [1]. The parameters *k*_cat_ (catalytic rate) and *K*_M_ (Michaelis-Menten constant) are regarded as steady-state parameters, even though the latter is attained at substrate concentration at half the maximum velocity of catalysis, which is far from pre-steady-state, closer to steady-state, and next to the zero-order zone if the substrate concentration ([S_0_]) at zero time is saturating. To begin with, the following may be true:

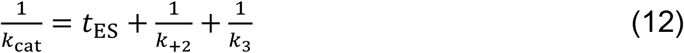

where *t*_ES_ is the duration for the formation of ES whose derivation is as described in the literature [19]. Equations (41c–44) of the paper [19] are being rederived to correct a technical error that was unfolded recently in the light of this research. This underpins the danger of abandoning a stepwise approach in derivations, for which regret and apology are expressed to the scientific community. However, the equation [19] from which *t*_ES_ (=1/*k*_ES_) is calculated does not present any technical problem of a derivational nature.

From Eq. (12) is the following:

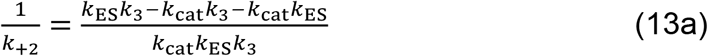

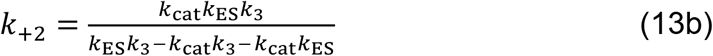

At this juncture, one can point out that if [*S*_0_] is much greater than [*E*_0_] and the duration of the assay (*τ*) is very short or rather transient, exp. (−*kτ*) [*S*_0_] (*k* is the pseudo-first-order rate constant for the utilization of substrate) should be approximately equal to [*S*_0_]. The duration of assay is the kind that is much less than the normal laboratory duration of assay. The time-scale of the assay in question is the micro-minute (or less). This time scale can be calculated for each value of [*E*_0_] for different substrate concentrations as described in the literature [19]. The implication is that the rate (*v*_1_) of formation of the enzyme-substrate complex (ES) is:

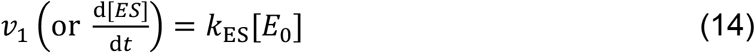

where a first-order rate constant, *k*_ES_ is approximately equal to *k*_1_[*S*_0_]. Equation (14) notwithstanding, a method for the determination of the maximum value of *k*_ES_ has been described in the literature [19]. The method for the determination of lower values of *k*_ES_ has been shown in the literature [11]. In order to derive the equation for *k*_-2_, one should recall Eq. (12), and following that, state first the equation of *k*_-1_ in terms of time as follows:

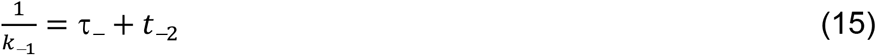

where *τ*_*−*_ (*i*.*e*.1/*k*_−11_) and *t*_−2_ are the durations of the dissociation of ES to free E and free S and the duration for the transformation or deactivation of E^#^S^#^ to ES respectively.

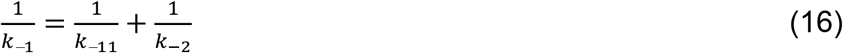

where *k*_*−*11_ is the first-order rate constant for the dissociation of ES to free E and free S. One can make *k*_−2_ subject of the formula in Eq. (16) to give:

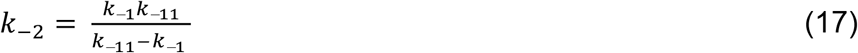

Given that *k*_−1_ = *K*_M_*k*_1_−*k*_cat_, the *K*_M_ can be expressed in terms of all the rate constants except *k*_+2_ and *k*_−1_ as follows: First *K*_M_*k*_1_−*k*_cat_ is substituted into Eq. (17) to give:

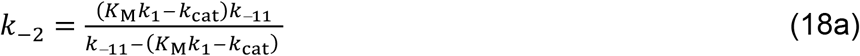

After rearrangement, *K*_M_ can be expressed as:

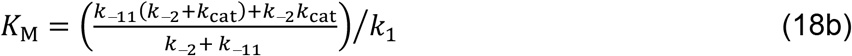

Following the example of Eq. (18b), and given that *k*_cat_ = *K*_M_ *k*_1_−*k*_−1_, the *K*_M_ can also be expressed in terms of all the rate constants except *k*_+2_ and *k*_cat_ as follows: First *K*_M_*k*_1_−*k*_−1_ is substituted into Eq. (13b) to give:

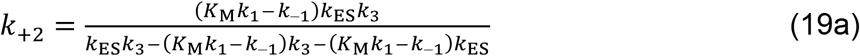

After rearrangement, *K*_M_ can be expressed as:

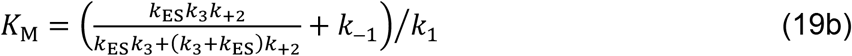

There is a need at this juncture to state that the process “E^#^S^#^ → EP” is the actual catalytic event, which can be given its exclusive duration without necessarily specifying the duration of E^#^S^#^ formation separately from the duration of the transition to EP.

It is not the characteristic of scientists to ignore what they consider trivial based on their conception of the simplicity of the mathematical content of an article, but to do so may lead to misinformation and ambiguity. Thus, while admitting that

Eqs (18b) and (19b) can give similar results, any difference should be the result of approximations of the original raw data. What should not be ignored are the equations, as corollaries, which express the mutual dependence of *k*_-1_ and *k*_cat_. Thus, bringing Eqs (18b) and (19b) together, simplifying, and making *k*_-1_ and *k*_cat_ subjects of the formula gives:

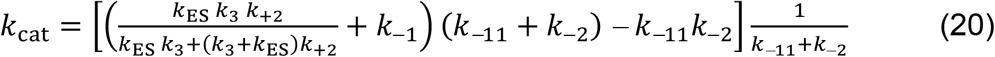

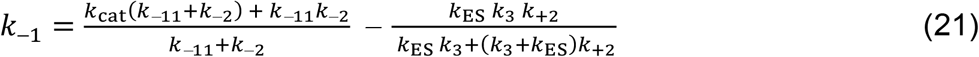

## 3. Results and Discussion

To begin with, it is necessary to point out that the result of this research with respect to usual Michaelian parameters may differ from those reported for the same enzyme in different concentrations and assay conditions in the literature [11, 19] for different reasons in accordance with the aims and objectives; comparison is taken to be unnecessary because the concerns covering the aims and objectives are quite different, with hardly any data for the first-order rate constants of the kind investigated in the literature. This research has, however, been able to produce equations for the determination of the values of hitherto unmeasurable first-order rate constants for the transition from ES to E^#^S^#^ and from the latter to ES. Similar to the view elsewhere [1], the process E^#^S^#^ → ES is possible, but the process EP → E^#^S^#^, let alone E + P → ES, is thermodynamically unfeasible. Upon algebraic appraisal and analysis of the equations reported in the literature [1], it was observed that the kinetic constants, *k*_cat_ and *K*_M_, cannot produce the values of the first-order rate constants, *k*_-2_ and *k*_+2,_ and if the latter values are known *ab initio* by whatever means, substituting them into the literature equations [1], cannot, beyond unnecessary subjectivity or sentiment, reproduce the elements of the Michaelis-Menten constant as evidenced in the outcome of algebraic appraisal and analysis (Eqs 6 and 11b), The equations (13b), (17b), (18b), and (19b) derived in this research can positively be used to quantify the corresponding parameter; quantification is, however, restricted to *k*_-2_ and *k*_+2_, though this requires information about the *K*_M_ and *V*_max_.

Proceeding further requires that certain points be made known. Since the velocity is linearly proportional to the substrate concentration if the latter is low, attributing *k*_cat_/*K*_M_ to a first-order zone is a little perplexing given that the constant, *k*_cat_, is asymptotically approached at much higher substrate concentration [17]. The admissibility of *K*_M_ as one best estimated in a mixed-order or pseudo-first-order zone calls into question the belief (or, mildly speaking, the suggestion) that the ratio *k*_cat_/*K*_M_ is best measured in a first-order zone. [17] If, as stated categorically, *K*_M_ is merely a ratio of the parameters (perhaps the sum of ratios given as (*k*_-1_/*k*_1_) + *k*_cat_/*k*_1_) estimated at the zero-order zone and the first-order zone, then it is an overt contradiction to describe *k*_cat_/*K*_M_ as a steady-state parameter [17]. This issue has been raised and discussed elsewhere. [20] So, in this study, zero-order kinetic parameters (specifically, the Michaelis-Menten parameters *K*_M_ and *V*_max_) are used rather than steady-state kinetic parameters, which is significant progress in this section.

As vividly shown in Table 1, the main Michaelian parameters are the zero-order parameters: 10.058 exp. (−4) M/min for the *V*_max_ and 60.527 g/L for the *K*_M_. The second-order rate constant for the formation of ES is well known in most general text books, but in the rate equation for the formation of ES, the first-order rate constant (preferably a pseudo-first-order rate constant) can be *k*_1_ [*E*_0_] or *k*_1_ [*S*_T_] depending on whether the assay was performed with a fixed concentration of the S and a varying concentration of the E or with a fixed concentration of the E and a varying concentration of the S. The maximum velocity *V*_max_ for product formation and release differs by 99.4% from the maximum velocity *V*_−max_ for ES dissociation into free E and free S; this is how *k*_-1_ compares to *k*_cat_. Zero-order dissociation constant is similar to the *K*_M_, the difference being 0.605 % of the *K*_M_.

**Table 1:**
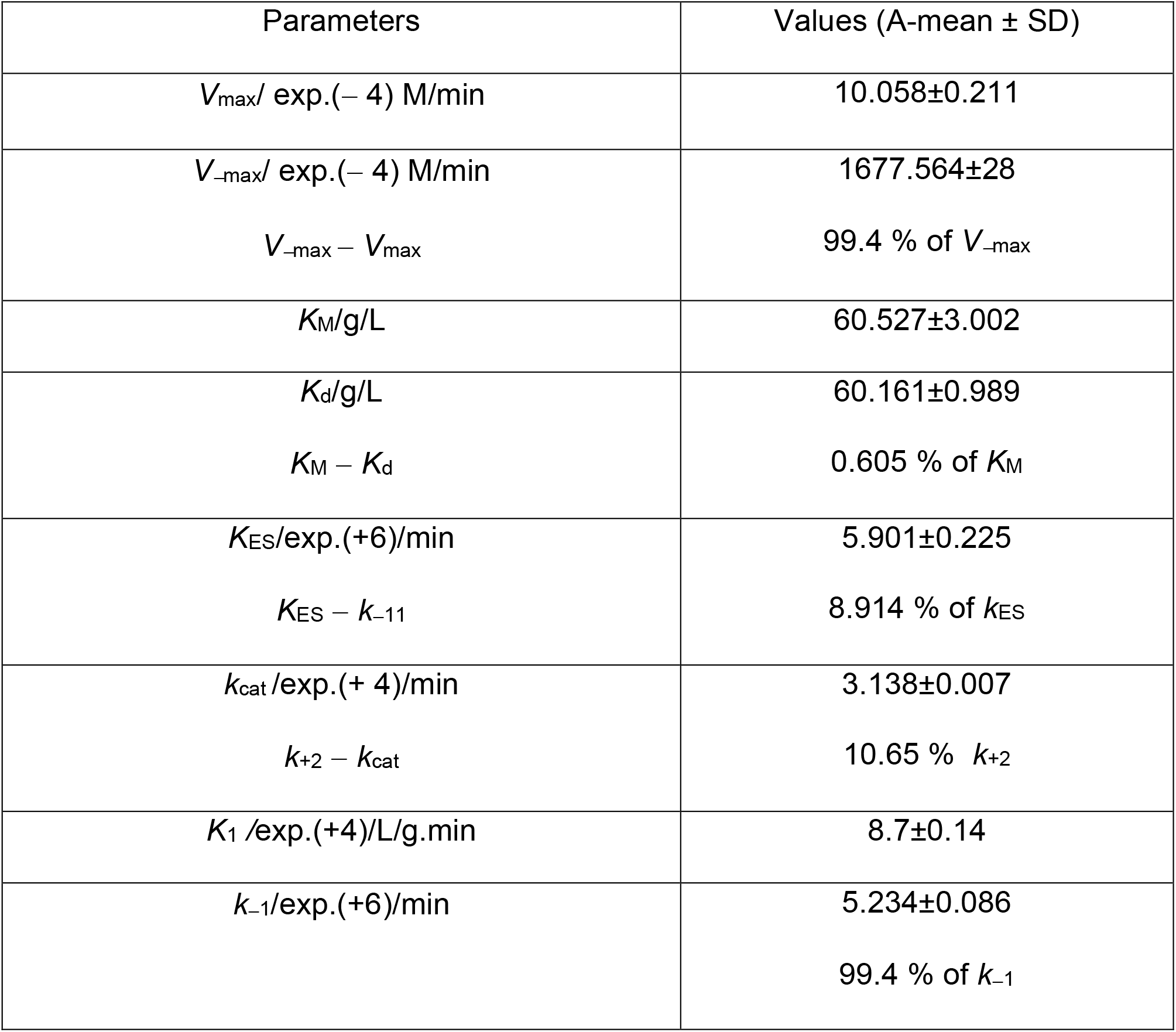
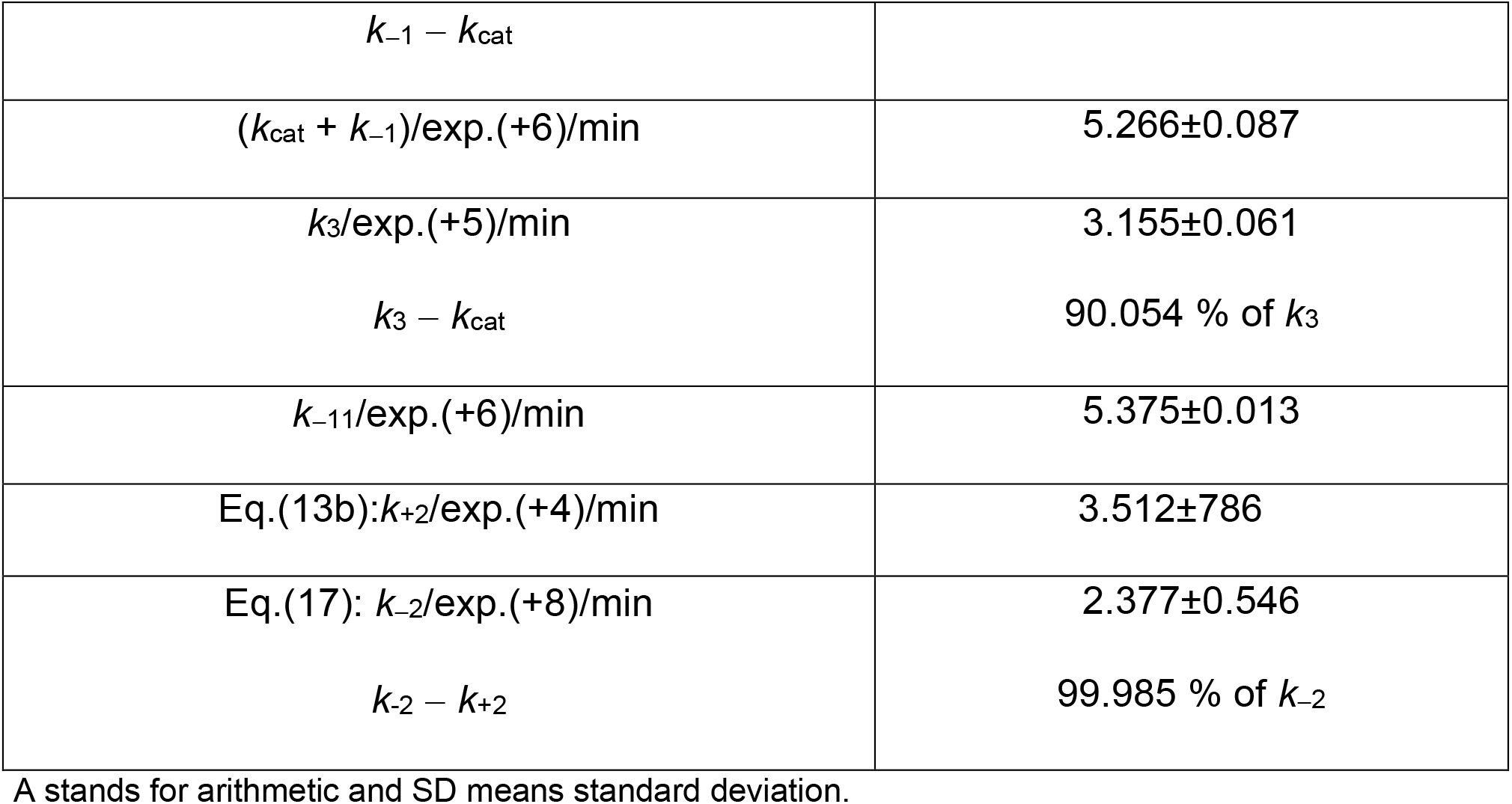
The values of kinetic parameters-the zero-order parameters

| Parameters | Values (A-mean ± SD) |
| --- | --- |
| $V_{\text{max}}/\text{exp.}(-4)\text{ M/min}$ | 10.058±0.211 |
| $V_{-\text{max}}/\text{exp.}(-4)\text{ M/min}$ | 1677.564±28 |
| $V_{-\text{max}} - V_{\text{max}}$ | 99.4 % of $V_{-\text{max}}$ |
| $K_M/\text{g/L}$ | 60.527±3.002 |
| $K_d/\text{g/L}$ | 60.161±0.989 |
| $K_M - K_d$ | 0.605 % of $K_M$ |
| $K_{ES}/\text{exp.}(+6)/\text{min}$ | 5.901±0.225 |
| $K_{ES} - k_{-11}$ | 8.914 % of $k_{ES}$ |
| $k_{\text{cat}}/\text{exp.}(+4)/\text{min}$ | 3.138±0.007 |
| $k_{+2} - k_{\text{cat}}$ | 10.65 % $k_{+2}$ |
| $K_1/\text{exp.}(+4)/\text{L/g.min}$ | 8.7±0.14 |
| $k_{-1}/\text{exp.}(+6)/\text{min}$ | 5.234±0.086 |
| | 99.4 % of $k_{-1}$ |
| $k_{-1} - k_{cat}$ | |
| $(k_{cat} + k_{-1})/\text{exp.}(+6)/\text{min}$ | $5.266 \pm 0.087$ |
| $k_3/\text{exp.}(+5)/\text{min}$ | $3.155 \pm 0.061$ |
| $k_3 - k_{cat}$ | 90.054 % of $k_3$ |
| $k_{-11}/\text{exp.}(+6)/\text{min}$ | $5.375 \pm 0.013$ |
| Eq.(13b): $k_{+2}/\text{exp.}(+4)/\text{min}$ | $3.512 \pm 786$ |
| Eq.(17): $k_{-2}/\text{exp.}(+8)/\text{min}$ | $2.377 \pm 0.546$ |
| $k_{-2} - k_{+2}$ | 99.985 % of $k_{-2}$ |

A stands for arithmetic and SD means standard deviation.

The first-order rate constants (*k*_cat_ and *k*_+2_) are similar in magnitude but differ, the difference being approximately 10.65 % of the *k*_+2_-value. The *k*_cat_ value differs from the *k*_3_ value by 90.054 % of the latter; the opposing processes, E + S → ES and ES **→** E + S, occurred at first-order rates, *k*_ES_ and *k*_*−*11_, respectively, which differ by approximately 8.914 % of the *k*_ES_. The *K*_M_ and *K*_d_ served the purpose of determining by calculation the values of *k*_3_ and *k*_*−*11_, respectively, as explained in the method section. As one should expect, *k*_-1_ is calculated by subtracting *k*_cat_ from the sum of *k*_-1_ and *k*_cat_ (Table 1), the equivalent of *K*_M_*k*_1_.

Next, one may consider matters of general interest connected to the issue of measurements, which this research addressed as part of its goal. It is already known that in any study on enzyme kinetics, there are measureable and unmeasurable quantities [17]. In this research, categorization is expanded to cover 1) primary measurables, 2) secondary measurables, and 3) tertiary measurables. Examples of the primary measurables are the mass concentrations of the substrate, enzymes, coenzymes, cofactors if applicable, and environmental conditions such as pH, ionic strength, temperature, and if necessary, pressure. Examples of the secondary measurables are the initial rates or the velocity of the catalytic action, changes in absorbance as applicable to conformational changes of the protein as enzyme or substrate, RNA, DNA, *etc*. Examples of tertiary measurables are those parameters that are either graphically or calculationally determined; such include Michaelis-Menten parameters, all first- and second-order rate constants, and the dissociation constant. Therefore, in this research, attention is paid to directly and indirectly determinable (measurable) rate constants in Michaelian enzyme-catalyzed reactions. The measurement in this research falls into the tertiary category. As shown in Table 1, they are first-order rate constants that the equations in the literature cannot adequately address; the rate constants are *k*_-2_, *k*_+2_, and *k*_3_.

Most importantly are the observed first-order rate constants that are greater than the usual rate constants, the *k*_cat_, and *k*_-1._ The implication as noted in the literature [1, 16, 20] is that the over-all duration for all the catalytic event, both physico-chemistry and biochemistry, is greater than the duration of any of the catalytic events. Each of the durations (not shown) is equal to the reciprocal of the zero-order, first-order rate constant. They are in the order of magnitude, the following: *k*_-2_>*k*_3_>*k*_+2_ (Table 1). The corresponding order of magnitude on a time-scale is: 1/*k*_−*2*_«1/*k*_3_<*k*_+2_.

The need for stability of enzyme-substrate complexes has been observed in the literature [1, 17, 21]. Following an encounter-complex formation, a kinetically driven process, with what has been described as the substrate’s right geometry in terms of structural and electrostatic orientations [1], the enzyme-substrate complex is formed: But this is purely governed by physico-chemical factors. The (bio) chemical factor is the function of the enzyme, which, through its internal mechanism, assumes an activated state that can also activate the substrate. This facilitates the breaking and making of bonds so that either a product or a substrate fragment or both can be released, depending on the nature of the substrate. Here the view of Van Slyke and Cullen [21] becomes very relevant, though they refer to “ordinary” rather than activated ES (E^#^S^#^). They admit that the process ES → E + S can occur because the interaction between S and E is noncovalent and dominated by hydrogen bonding and electrostatic effects, which are strongly subject to environmental perturbation; but upon assumption of the chemistry by the enzyme, a stronger binding interaction is promoted to enable the breaking and making of new bonds for product formation. If on the other hand a stronger binding interaction leads to rigid covalent bond formation without any form of structural flexibility, then the enzyme is said to be inhibited. All these point to the importance of catalytically functional activated complex. As shown in Table (1) the rate constants, *k*_−2_ and *k*_+2_, for the deactivation of E^#^S^#^ to ES and the converse respectively, are widely different, the difference being approximately 99.985 % of the *k*_−2_. This may account for the large value of *k*_−1_ which is exceedingly greater than *k*_cat_. This may imply that there is a very high probability (0.99985), *k*_−2_/(*k*_−2_+*k*_+2_), of the occurrence of occasional deactivation of the activated ES complex.

It behoves the process, chemical, engineers *etc*. to continue to consider the following fundamental facts: the role of thermodynamic temperature as a driver of kinetic aspects of enzyme-catalyzed reactions beyond kinetic stability; this implies that the speed at which an encounter-complex is formed depends on the translational diffusion coefficient, and consequently the rate of effective collision and encounter-complex formation [22, 23]. Therefore, any factor—inorganic or organic osmolyte, genetic modification, or perhaps immobilization—that can sustain the stability of the enzyme while increasing the temperature even above the optimum temperature should be encouraged. In this era of the need for environmentally friendly fuel consumption, the optimization of the production of bio-fuels requires that the *k*_+2_ values be boosted while diminishing the *k*_-2_ values. The concern of clinical dieticians, the pharmaceutical industry, and medics is how best to compulsorily control diabetics; in this regard, ways of boosting the *k*_-2_ values should be encouraged.

## 4. Conclusion

All the equations for the calculation of zero-order first-order rate constants (ZOFORC) for the activation of the enzyme-substrate (ES) complex and its deactivation, *k*_+2_ and *k*_-2_ respectively, were derived. The value of ZOFORC for the dissociation of the enzyme-product (EP) complex to free E and P is 3.155 exp. (+5)/min; the values of *k*_+2_ and *k*_-2_ are 3.513 exp.(+4) and 2.377 exp. (+8)/min, respectively. Ultimately, it is imperative for all stakeholder groups, including dieticians, medics, paramedics, technologists, engineers, *etc*., to devise means of controlling the enzymatic rate of catalysis by manipulating the magnitudes of *k*_+2_ and *k*_-2_ in particular. The derived equations can be fitted to the experimentally generated and calculated data. Since this research was carried out without regard to the enzyme’s optimal conditions, future research should entail conducting assays under optimum conditions so as to verify possible variations in the ZOFORC values when compared with values generated outside optimum conditions.

## 5. Experimental

### 5.1. MATERIALS AND METHODS

#### 3.1.1 Materials

##### 3.1.1.1 Chemicals

The enzyme which was assayed is *Aspergillus oryzae* alpha-amylase (EC 3.2.1.1) and, insoluble potato starch, was the substrate; both were purchased from Sigma–Aldrich, USA. Tris 3, 5—di-nitro-salicylic acid, maltose, and sodium potassium tartrate tetrahydrate were purchased from Kem Light Laboratories in Mumbai, India. Hydrochloric acid, sodium hydroxide, and sodium chloride were purchased from BDH Chemical Ltd., Poole, England. Distilled water was purchased from the local market. The molar mass of the enzyme is approximately 52 k Da [24, 25]. Distilled water was purchased from the local market. As a word of caution, readers of this paper should be aware that the use of the same enzyme in articles by the same author(s) is strictly due to budgetary constraints; however, this is not a serious concern because each paper addresses different issues, such as the evaluation of new models.

##### 3.1.1.2 Equipment

An electronic weighing machine was purchased from Wensar Weighing Scale Limited, and a 721/722 visible spectrophotometer was purchased from Spectrum Instruments, China; a *p*H meter was purchased from Hanna Instruments, Italy.

### 5.2 Methods

#### 5.2.1 Preparation of reagents and assay

The method of assaying the enzyme is Benfield’s method [26], which uses gelatinized potato starch, whose concentration range is 5–10 g/L. The reducing sugar produced upon hydrolysis of the substrate at 20 °C using maltose as a standard was determined at 540 nm with an extinction coefficient approximately equal to 181 L/mol.cm. The assay took 3 minutes to complete. A mass concentration equal to 1.667 mg/L of *Aspergillus oryzae* alpha-amylase was prepared in a Tris-HCl buffer at *p*H 6.9; there were special considerations in the choice of *p*H and temperature. The evaluation of new equations was the only overriding interest.

#### 5.2.2. The determination of rate constants

The pseudo-first-order rate constant for the utilization of gelatinized starch is determined as described in the literature [27], while the second-order rate constant for the formation of the enzyme-substrate (ES) complex and the duration of its formation are determined as described elsewhere [19]. While *V*_max_(*k*_cat_ [*E*_0_]) is a well-known parameter, *V*_-max_(*k*_-1_[*E*_0_]) for the release of free E and S from ES is not well-known. The first-order rate constant, *k*_−1_, for the release of free E and free S is also determined as described in the literature [19]. The Lineweaver-Burk [28] plot was used for the determination of the *K*_M_ and *V*_max_. Equations (13b) and (17) were for the calculation of *k*_+2_ and *k*_-2_ respectively. The determination of *k*_3_ is as described in the literature [19], but a corrected version is stated herein as follows:

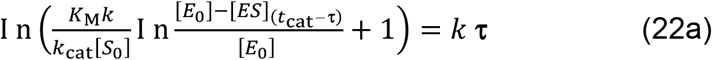

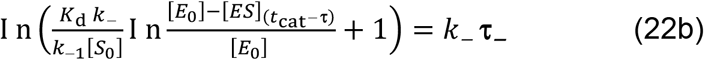

To refresh your memory, the rate constants *k*_cat_ and *k*_-1_ represent the catalytic first-order rates for the processes ES → E^#^S^#^→EP→E+P and E^#^S^#^ → ES→E + S, respectively. The durations *τ* and *τ*_-_ are the durations for the processes leading to the release of product and free enzyme and free enzyme and substrate, respectively.

#### 5.3 Statistical analysis

Assays were conducted in triplicates. Micro-Soft Excel was used for the determination of the standard deviation (SD) for the arithmetic mean values.

## Acknowledgements

The management of the Royal Court Yard Hotel, Agbor, Delta State, Nigeria, is thanked for the supply of electricity during the preparation of the manuscript. The provider of the QuillBot grammar checker is thanked for the proofreading services.

